# Divergent landscapes of positive and negative selection signatures across residue-resolved human-virus protein-protein interaction interfaces

**DOI:** 10.64898/2026.04.10.717550

**Authors:** Wan-Chun Su, Yu Xia

## Abstract

Virus-targeted host proteins evolve under dual selective pressures. Negative selection preserves within-host interactions, while positive selection promotes adaptive changes to evade viral engagement. Viral and endogenous within-host partners can compete for binding, bringing distinct pressures together on the same interaction interface. Yet, the spatial organization of distinct selective pressures across virus-targeted host proteins, and how such pressures manifest across diverse interaction contexts, remains largely unknown. Here, we integrate an evolutionarily annotated map of human-virus protein-protein interactions (PPIs) with intra-protein residue-residue contact maps to probe the spatial organization of residue-level selective pressures across PPI interfaces of virus-targeted host proteins. Across all PPI interfaces collectively, we find that residues under positive selection are spatially clustered, whereas those under negative selection are broadly dispersed, with additional spatial segregation between positive and strongly negatively selected sites. Moreover, while positive selection is unevenly distributed across interfaces bound exclusively by viral proteins (exogenous-specific), they are more uniformly distributed across interfaces shared between viral and within-host partners (mimic-targeted), suggesting that adaptive pressure from viral targeting acts on the entire mimic-targeted interface, whereas it acts on only a subset of the exogenous-specific interface. Strikingly, clustering of positively selected residues is more pronounced between mimic-targeted and other interface types than within exogenous-or endogenous-specific interfaces alone, suggesting that mimic-targeted interfaces may serve as focal points of adaptive evolution. Overall, our multiscale framework of PPI interfaces and residue-level contacts reveals heterogeneous, context-dependent landscapes of selective pressures across virus-targeted host proteins, providing a high-resolution view of how adaptation and constraint are intricately balanced and coordinated within the host.

## Introduction

Protein evolution is shaped by a combination of selective pressures. Positive selection promotes adaptive changes, whereas negative selection constrains variation to preserve structural and functional integrity [1–5]. Across the proteome, these forces often act in complementary ways: positively selected residues tend to occur on the surface of a protein, while negatively selected residues are more frequently buried within the core of a protein [5–9]. Protein-protein interaction (PPI) interfaces are particularly important regions, as they occupy a position between the protein core and surface [10–12], where residues are exposed enough to participate in binding while remaining constrained to preserve structural stability. PPI interfaces are especially relevant for proteins engaged in diverse interactions, such as virus-targeted host proteins, where adaptive responses to exogenous pressures act alongside constraints to maintain endogenous functions.

Prior studies have detected residues under positive selection [13–16] and demonstrated the coexistence of positive and negative selection [17–22] on individual virus-targeted host proteins. Notably, positively selected residues were found to be located at virus-targeted PPI interfaces of host proteins [14,16,20–24]. While the presence of positively selected residues at virus-targeted host proteins, particularly at their PPI interfaces, is well established, quantitative structural characterization of how positive selection is spatially organized across these interfaces remains limited. Across mammalian proteins more broadly, positive selection has been found to cluster near active sites on metabolic enzymes or at immune-related binding interfaces [25]. Such studies typically assess clustering relative to the entire protein structure rather than within individual interfaces, and thus it remains unclear whether positively selected residues tend to be distributed across an entire interface (i.e., whether an interface as a whole constitutes a cluster) or are instead concentrated within discrete subregions of an interface. By examining how positive selection is arranged within PPI interfaces themselves, rather than relative to the whole protein structure, we can gain a higher-resolution view of how adaptive changes are distributed across functional regions, potentially revealing patterns of evolutionary coordination that may otherwise be obscured at the protein-wide scale.

Virus-targeted host proteins as a whole are generally under strong negative selection, reflecting the tendency for viruses to target conserved host proteins [23,26,27]. Yet, within host PPI interfaces, adaptive changes can still occur, raising questions about how positive and negative selection are arranged relative to one another across the same interface. Structural and biochemical studies have shown that functionally important residues are generally conserved under negative selection [28–31], but these constraints are not always confined to functional sites. For example, while enzyme active sites are under strong negative selection, such evolutionary constraints can propagate across regions beyond those functional sites [32]. Hence, evolutionarily constrained regions do not necessarily align with functionally defined regions, such as interaction interfaces. It remains unclear whether positively selected residues are strictly limited to viral-binding sites, or whether they may also occur in neighboring sites, as has been observed for negative selection. Together, these considerations underscore the importance of spatially resolving distinct selective pressures across PPI interfaces to better understand the interplay between evolutionary constraint and adaptation in virus-targeted host proteins.

PPI interfaces on virus-targeted host proteins experience diverse interaction contexts: some regions are bound exclusively by viral proteins (exogenous-specific interfaces), others only by within-host proteins (endogenous-specific interfaces), and some are shared and competitively bound by viral and within-host proteins (mimic-targeted interfaces). Recent analyses of host PPI interface evolution have revealed general trends in how viral targeting shapes host evolution, through comparisons of average evolutionary rates across entire interfaces [23,24]. However, such interface-level averaging obscures finer-scale, residue-level patterns of positive and negative selection. Importantly, at mimic-targeted interfaces, the pressures to maintain endogenous binding and to evade viral engagement occur within the same interface, likely giving rise to more complex residue-level evolutionary landscapes than interfaces exclusive to viral or within-host partners. The spatial organization of selective pressures across these diverse interaction contexts remains largely uncharacterized, highlighting the need for analyses that resolve selection at single-residue resolution while accounting for the specific interaction context of each residue.

In this work, using an established human-virus structural interaction network with residue-level evolutionary rates [24], we constructed an evolutionary rate-informed residue-residue contact map that captures residue-level interactions between selective pressures on host PPI interfaces. By comparing this contact map to randomized maps generated through residue-label permutations, we assess the spatial organization of selective pressures across the PPI interfaces of virus-targeted host proteins. At the global scale (across all PPI interfaces collectively), we find that positively selected residues are highly clustered, whereas negatively selected residues are more uniformly distributed. Positively selected residues are also spatially segregated from strongly negatively selected residues, a pattern that cannot be explained by the spatial clustering of positively selected residues. Furthermore, within virus-targeted interfaces, while positively selected residues are unevenly distributed across exogenous-specific interfaces, they are more uniformly distributed across mimic-targeted interfaces, suggesting that exogenous-specific regions contribute partially, whereas mimic-targeted regions are fully involved in the host-virus evolutionary arms race. Strikingly, cross-interface clustering of positively selected residues between mimic-targeted and exogenous-or endogenous-specific interfaces exceeds clustering within exogenous-or endogenous-specific interfaces alone, suggesting that mimic-targeted interfaces may act as focal points of adaptive pressure, potentially influencing adaptive changes at neighboring interfaces. Taken together, our findings provide quantitative evidence for partitioning across PPI interfaces on virus-targeted host proteins, both between different types of selective pressures and across different interaction contexts, reflecting a coordinated, structured balance between adaptive flexibility and constraint within the host.

## Results

### Residue-residue contact graph of all interfacial residues within the human-virus structural interaction network

We constructed a residue-residue contact graph of all virus-human and within-human interfacial residues on virus-targeted human proteins (hereafter referred to as human target proteins). These interfacial residues, along with their site-specific evolutionary rates (dN/dS), were originally generated in our recently developed human-virus structural interaction network [24], which mapped existing virus-human and within-human PPIs to experimentally-determined three-dimensional (3D) structures from the Protein Data Bank [33] or homology-based structural templates (see **Material and Methods** for more details). While the previous work focused on interface-level summaries (e.g., average site-specific dN/dS per interface or fractions of residues above a certain dN/dS threshold), the present study represents the first residue-level analysis of these interfacial residues, retaining site-specific dN/dS values and treating each residue as an individual unit rather than aggregating rates across an interface.

In the residue-residue contact graph (**Figure 1A**), each node represents an interfacial residue, each edge depicts two interfacial residues in contact (minimum atomic distance between residues < 5 Å), and each cluster corresponds to an individual human target protein (see **Materials and Methods** for more details on contact graph construction). Interfacial residues (nodes) were categorized based on their site-specific dN/dS values (residue-level evolutionary rates): (i) positively selected (“Pos”, dN/dS > 1), (ii) strongly negatively selected (“SNeg”, dN/dS < 0.1), and (iii) all others (“Other”, 0.1 ≤ dN/dS ≤ 1). Here, we used a more stringent negative selection threshold (dN/dS < 0.1), rather than the conventional dN/dS < 1 threshold, to ensure a more balanced comparison with positive selection (dN/dS > 1). Residue-residue contacts (edges) were categorized based on the contact type: (i) “Pos-Pos”: two positively selected residues in contact, (ii) “SNeg-SNeg”: two strongly negatively selected residues in contact, (iii) “Pos-SNeg”: a positively selected residue and a strongly negatively selected residue in contact, and (iv) “Other”: all other pairs of residues in contact. The number of proteins and contact counts are shown in **Table S1**.

**Figure 1.**
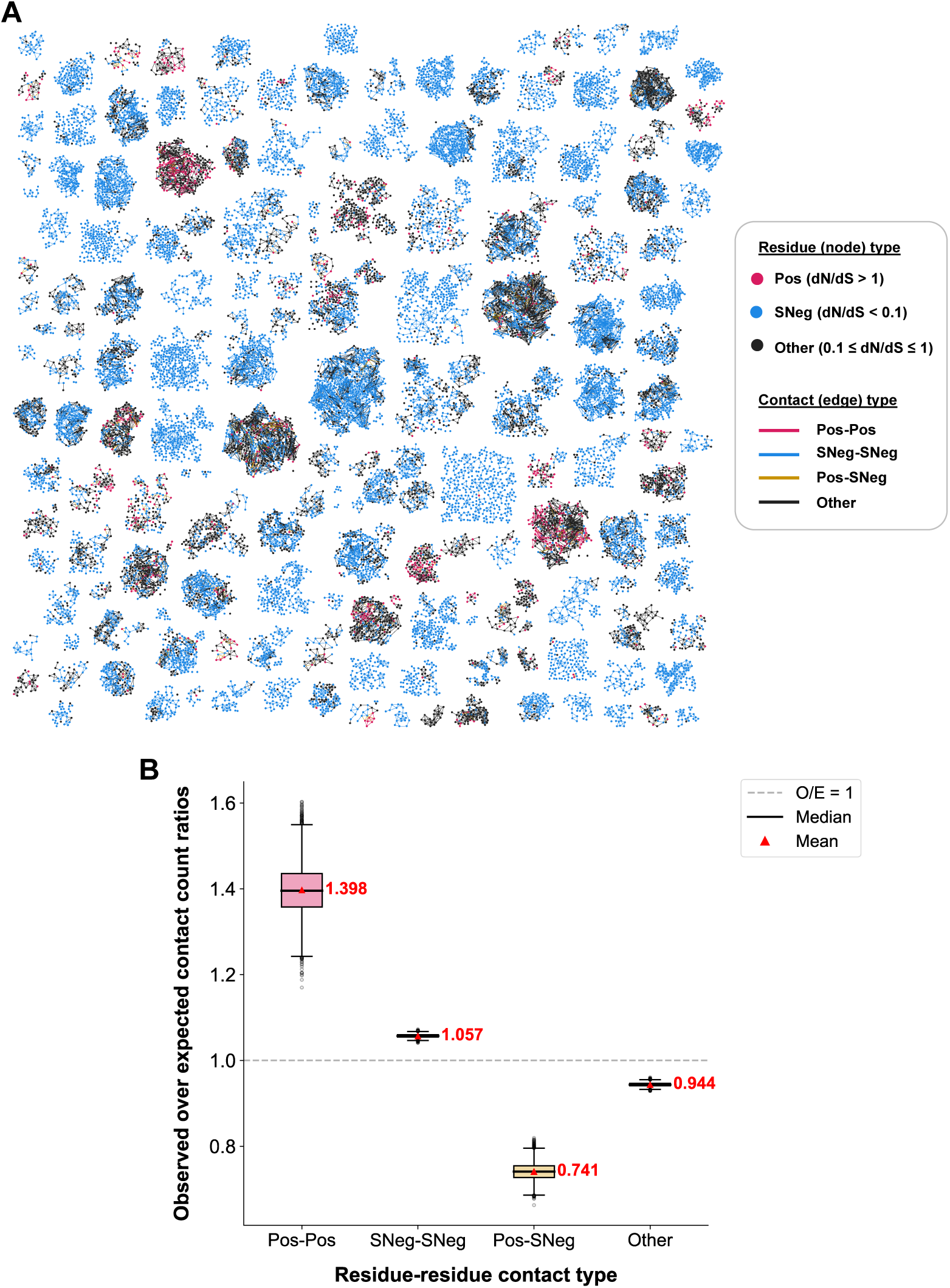
Residue-residue contact graph of all interfacial residues and observed-over-expected contact count ratios for different contact types. **(A)** Residue-residue contact graph of all interfacial residues on human target proteins (virus-targeted human proteins) in the human-virus structural interaction network, visualized using Graphviz [63]. Each cluster in the contact graph represents an individual human target protein. Nodes represent interfacial residues on human target proteins and edges represent a pair of interfacial residues (on the same protein) in contact. Nodes and edges in the contact graph are labeled according to their corresponding dN/dS value and contact type, respectively. Nodes are classified as (i) “Pos” (dN/dS > 1), (ii) “SNeg” (dN/dS < 0.1), or (iii) “Other” (0.1 ≤ dN/dS ≤ 1). We note that a more stringent threshold for negative selection (dN/dS < 0.1), rather than the conventional dN/dS < 1 threshold, was used to enable a more balanced comparison with positive selection (dN/dS > 1). Edges are classified as: (i) “Pos-Pos” (two positively selected residues in contact), (ii) “SNeg-SNeg” (two strongly negatively selected residues in contact), (iii) “Pos-SNeg” (a positively selected residue and a strongly negatively selected residue in contact), or (iv) “Other” (all other pairs of residues in contact, which contains three subcategories: “Pos-Other”, “SNeg-Other”, and “Other-Other”). **(B)** Boxplots depicting the distribution of observed-over-expected (O/E) contact counts for each residue-residue contact type across 10,000 randomized (expected) contact graphs. In each randomization trial, the observed number of contacts remains unchanged, while the expected number is calculated from a randomized contact graph generated by permuting node labels within each protein. The mean and median O/E across the randomization trials are denoted by a red triangle and a black horizontal line, respectively. We use the mean O/E (labeled in red beside the corresponding distribution) as the overall O/E ratio for a contact type. We define enrichment as the extent to which the overall O/E ratio deviates from 1 (baseline expectation, annotated as a gray dashed line). As all overall O/E ratios show significant deviation from 1 (*p* < 0.001, two-tailed one-sample t-test against 1), we focus our analysis on effect size rather than statistical significance.

### Positively selected and strongly negatively selected interfacial residues exhibit distinct spatial organization

To examine the spatial organization of different selective pressures (“Pos”, “SNeg”, and “Other”) within the residue-residue contact graph, we first computed observed-over-expected (O/E) contact count ratios for each contact type (“Pos-Pos”, “SNeg-SNeg”, “Pos-SNeg”, and “Other”) across 10,000 within-protein node-label randomization trials (see **Materials and Methods** for more details on computing O/E ratios). For each contact type, we used the mean O/E ratio from the 10,000 randomization trials as the representative (overall) O/E ratio for that contact type. The overall O/E ratio for a contact type was assessed as follows: (i) overall O/E >> 1: enrichment of this contact type, (ii) overall O/E ≈ 1: little to no enrichment or depletion of this contact type, (iii) overall O/E << 1: depletion of this contact type. We note that effect sizes were used as the primary measure of enrichment rather than statistical significance, as all overall O/E ratios are significantly different from 1 (*p* < 0.001, two-tailed one-sample t-test against 1).

Figure 1B illustrates the distribution of O/E ratios (based on 10,000 randomization trials) for each contact type across the contact graph, with overall O/E ratios labeled in red. We observe high enrichment of Pos-Pos contacts (overall O/E for Pos-Pos = 1.398 >> 1) and very little to no enrichment of SNeg-SNeg contacts (overall O/E for SNeg-SNeg = 1.057 ≈ 1), indicating that while positively selected residues are spatially clustered, strongly negatively selected residues are more uniformly distributed on PPI interfaces of human target proteins. Such trends remain qualitatively consistent on a per-protein basis (**Figure S1**; see **Materials and Methods** for details on calculating per-protein contact count enrichments), and for different residue-residue contact thresholds (**Figure S2**).

### Pronounced spatial segregation between positively selected and strongly negatively selected interfacial residues

In addition to exhibiting distinct spatial organization across PPI interfaces of human target proteins, positively and strongly negatively selected residues also demonstrate substantial segregation from one another, as shown by the large depletion in Pos-SNeg contacts in the residue-residue contact graph (overall O/E for Pos-SNeg = 0.741 << 1, Figure 1B). Together, these results are consistent with a non-random, spatially structured organization of positive and strong negative selection within PPI interfaces of human target proteins.

Proteins with low numbers of Pos or SNeg residues may potentially introduce noise and reduce the reliability of the contact graph-wide Pos-SNeg O/E ratio. Hence, we created subgraphs of our original contact graph where we isolated proteins with sufficient numbers of Pos and SNeg residues (i.e., at least *n* Pos and *n* SNeg interfacial residues, where *n* = 2, 3, 4, or 5) (**Figure S3**). Even after excluding proteins with low Pos or low SNeg counts, the overall O/E ratio for Pos-SNeg contacts consistently remains below 1, demonstrating the robustness of the spatial separation between Pos and SNeg residues. Similarly, the enrichment and depletion trends for other contact types remain consistent.

The observed spatial separation between Pos and SNeg residues may simply be a by-product of Pos clustering, rather than reflecting genuine spatial segregation between Pos and SNeg residues. To test this hypothesis, we created subgraphs of our original contact graph where we removed proteins with varying degrees of within-protein Pos clustering (i.e., per-protein O/E for Pos-Pos contacts > *t*, where *t* = 1.45, 1.35, 1.25, 1.15, and 1.05). We re-evaluated the overall O/E ratios for Pos-SNeg contacts within these subgraphs and found that Pos-SNeg contacts remain depleted to a similar degree regardless of varying degrees of within-protein Pos clustering (**Figure S4**), demonstrating that the observed spatial separation between positively and strongly negatively selected residues is not driven by clustering of positively selected residues.

### Spatial selection patterns of all interfacial residues are largely driven by interactions involving positively selected residues

Residue-residue contacts that involve at least one “Other” (neither positively selected nor strongly negatively selected) residue show little to no enrichment or depletion in the contact graph (overall O/E for “Other” contacts = 0.944 ≈ 1, Figure 1B). Even when these contacts are divided into three subcategories (“Pos-Other”, “SNeg-Other”, and “Other-Other”), the corresponding effect sizes for the O/E contact count ratios in each subcategory remain small (**Figure S5**). By contrast, contacts between positively selected residues and between positively and strongly negatively selected residues exhibit the largest and second largest effect sizes, respectively (Pos-Pos overall O/E = 1.398 >> 1 and Pos-SNeg overall O/E = 0.741 << 1, Figure 1B). These results are especially striking given that positively selected residues account for a tiny proportion (5%) of the entire set of interfacial residues in the contact graph. Such findings suggest that the spatial arrangement of selective pressures across PPI interfaces of human target proteins are predominantly driven by positively selected residues, with strongly negatively selected residues contributing to a lesser yet still appreciable extent. Moreover, these results suggest that any small effects seen for “Other” contacts likely reflect secondary effects arising from interactions involving positively and strongly negatively residues.

### Case studies

Here, we present two representative examples highlighting the distinct spatial arrangements of positively and negatively selected residues across PPI interfaces of human target proteins (Figure 2). Figure 2A illustrates the human tumor necrosis factor-related apoptosis-inducing ligand receptor 2 (TRAIL-R2; semi-transparent light gray surface) in complex with its endogenous human partner, tumor necrosis factor-related apoptosis inducing ligand (TRAIL; blue ribbon), superimposed with its interaction with an exogenous viral partner, UL141 (from human cytomegalovirus; orange ribbon). TRAIL-R2, in complex with its ligand TRAIL, mediates programmed cell death in infected cells to limit further infection, while UL141 interferes with this process by competing with TRAIL for binding to TRAIL-R2 [34–39]. Figure 2B displays the surface of TRAIL-R2, where the entire set of interfacial residues mediating the two interactions are highlighted and colored based on corresponding residue-specific selective pressure labels: red, positively selected (“Pos”); blue, strongly negatively selected (“SNeg”); and dark gray, all other interfacial residues. Non-interfacial residues are unhighlighted (semi-transparent light gray). We found clustering of Pos residues near one end of TRAIL-R2 and a more uniform distribution of both SNeg and all other interfacial residues. In particular, the two Pos residues located at the tip of TRAIL-R2 (E78 and D109) have been shown to be critical for UL141 binding to TRAIL-R2, as double mutations on these two residues resulted in a tenfold decrease in UL141 binding affinity [37], suggesting that the observed cluster of Pos residues represents the host’s localized strategy for evading viral binding. All the interfacial residues lie within the three cysteine-rich domains (CRDs) of TRAIL-R2: CRD-1 (residues E78–C94), CRD-2 (residues I95–C137), and CRD-3 (residues Q138–C178), where CRD-2 and CRD-3 mediate interactions with TRAIL [37,40]. Hence, the broad dispersion of both SNeg and all other interfacial residues across these domains likely reflects the host’s mechanism for maintaining endogenous binding to TRAIL.

**Figure 2.**
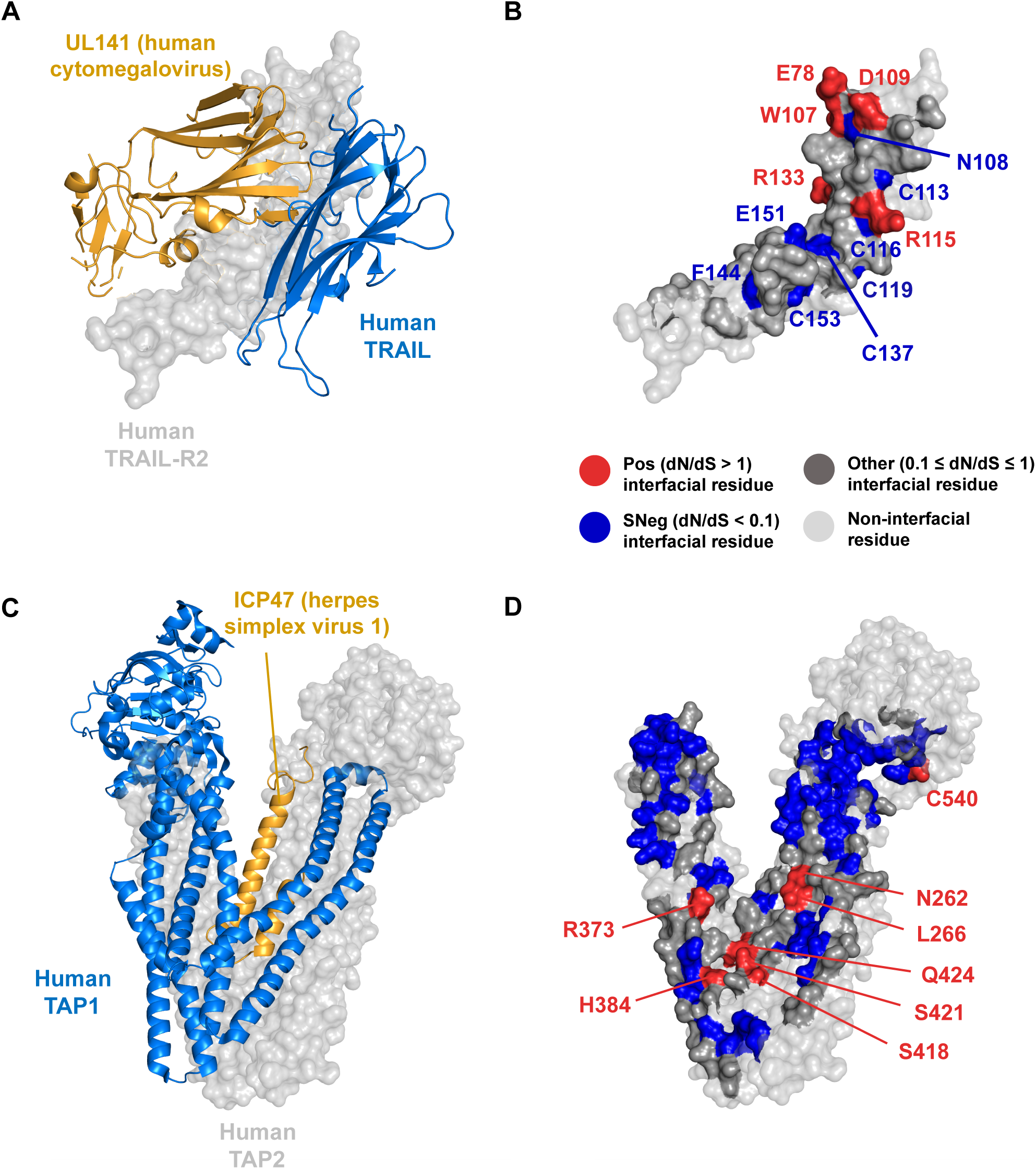
Case studies illustrating distinct spatial arrangements of positively and negatively selected interfacial residues on human target proteins. **(A)** Human target protein, TRAIL-R2 (light gray semi-transparent surface, center), in complex with human cytomegalovirus protein UL141 (orange ribbon, left) overlaid with the interaction between TRAIL-R2 and human TRAIL (blue ribbon, right), represented by PDB: 4i9x [37] and PDB: 1du3 [34]. **(B)** The surface of TRAIL-R2 with interfacial residues highlighted and colored based on their corresponding residue-specific dN/dS values: red depicts positively selected interfacial residues (Pos), blue depicts strongly negatively selected interfacial residues (SNeg), and dark gray depicts all other interfacial residues. Light gray (semi-transparent) depicts non-interfacial residues. Both Pos and SNeg residues are labeled with their one-letter amino acid code followed by the corresponding residue position on TRAIL-R2. Pos residues are clustered at one end of TRAIL-R2, while SNeg and all other interfacial residues are more uniformly distributed across the interface. **(C)** Human target protein, TAP2 (light gray semi-transparent surface), in complex with herpes simplex virus 1 protein ICP47 (orange ribbon) and its endogenous partner TAP1 (blue ribbon), represented by PDB: 5u1d [45]. **(D)** The surface of TAP2 with interfacial residues highlighted and colored based on their corresponding residue-specific dN/dS values: red depicts positively selected interfacial residues (Pos), blue depicts strongly negatively selected interfacial residues (SNeg), and dark gray depicts all other interfacial residues. Light gray (semi-transparent) depicts non-interfacial residues. Pos residues are labeled with their one-letter amino acid code followed by the corresponding residue position on TAP2. Pos residues are clustered at the inner cavity of TAP2. SNeg and all other interfacial residues are more uniformly distributed across the interface, with apparent spatial segregation between SNeg and Pos residues around the cluster of Pos residues. 3D protein complex structure images were created using PyMOL [64].

Figure 2C illustrates the human transporter associated with antigen processing 2 (TAP2; semi-transparent light gray surface) in complex with its endogenous human partner, transporter associated with antigen processing 1 (TAP1; blue ribbon), and an exogenous viral partner, ICP47 (from herpes simplex virus 1; orange ribbon). In the host, TAP2 and TAP1 together form the TAP complex, which transports peptides into the ER for downstream antigen presentation [41–43]. ICP47 inhibits this process by binding TAP, primarily at TAP2 [44,45]. Figure 2D displays the surface of TAP2, where the entire set of interfacial residues mediating the two interactions are highlighted and colored based on corresponding site-specific selective pressure labels, as in the above example. Non-interfacial residues are unhighlighted (semi-transparent light gray). We found clustering of Pos residues near the inner cavity of TAP2. Indeed, previous studies have shown that ICP47-TAP interactions are largely mediated by contacts between the loop connecting the two helices of ICP47 (lower region of ICP47, depicted in Figure 2C) and the inner cavity of TAP [45]. In particular, interfacial residues H384 and S421 on TAP2 (annotated in Figure 2D) were shown to interact with residues on ICP47 that are critical for binding and inhibiting the TAP complex [45,46]. In contrast to the strong clustering of positively selected interfacial residues, non-positively selected interfacial residues (both SNeg and all other interfacial residues) are more uniformly distributed across the entire interface (Figure 2D) and roughly correspond to the regions bound by TAP1 (Figure 2C). SNeg residues appear to be additionally segregated from Pos residues, as shown by the sparser distribution of SNeg residues near the cavity of TAP2, where Pos residues are clustered (Figure 2D). The observed clustering of positive selection and segregation between positive and strong negative selection across interfacial residues on TAP2 likely reflects the host’s strategy to separate localized virus-driven adaptation from more widespread functional constraints that preserve the TAP1-TAP2 complex.

### Within virus-targeted interfaces, spatial distribution of positively selected residues distinguishes exogenous-specific and mimic-targeted sub-interfaces

To determine whether different types of interfaces on human target proteins exhibit distinct residue-residue contact enrichment patterns, we generated subgraphs derived from the original contact graph, with each subgraph containing only the interfacial residues and residue-residue contacts corresponding to a specific interface category: (i) the entire virus-targeted (all exogenous) interface (**Figure S6**), and (ii) the entire within-human (all endogenous) interface (**Figure S7**). We further split these two interface contact graphs into three sub-interface contact graphs (Figure 3): (i) exogenous-specific sub-interface (targeted only by viral proteins), (ii) mimic-targeted sub-interface (targeted by both viral and human proteins), and (iii) endogenous-specific sub-interface (targeted only by human proteins). For each interface and sub-interface contact graph, we used the same procedure as for the original contact graph to assess enrichment or depletion of a contact type. The number of proteins and contact counts for each interface-specific contact graph are shown in **Table S1**.

**Figure 3.**
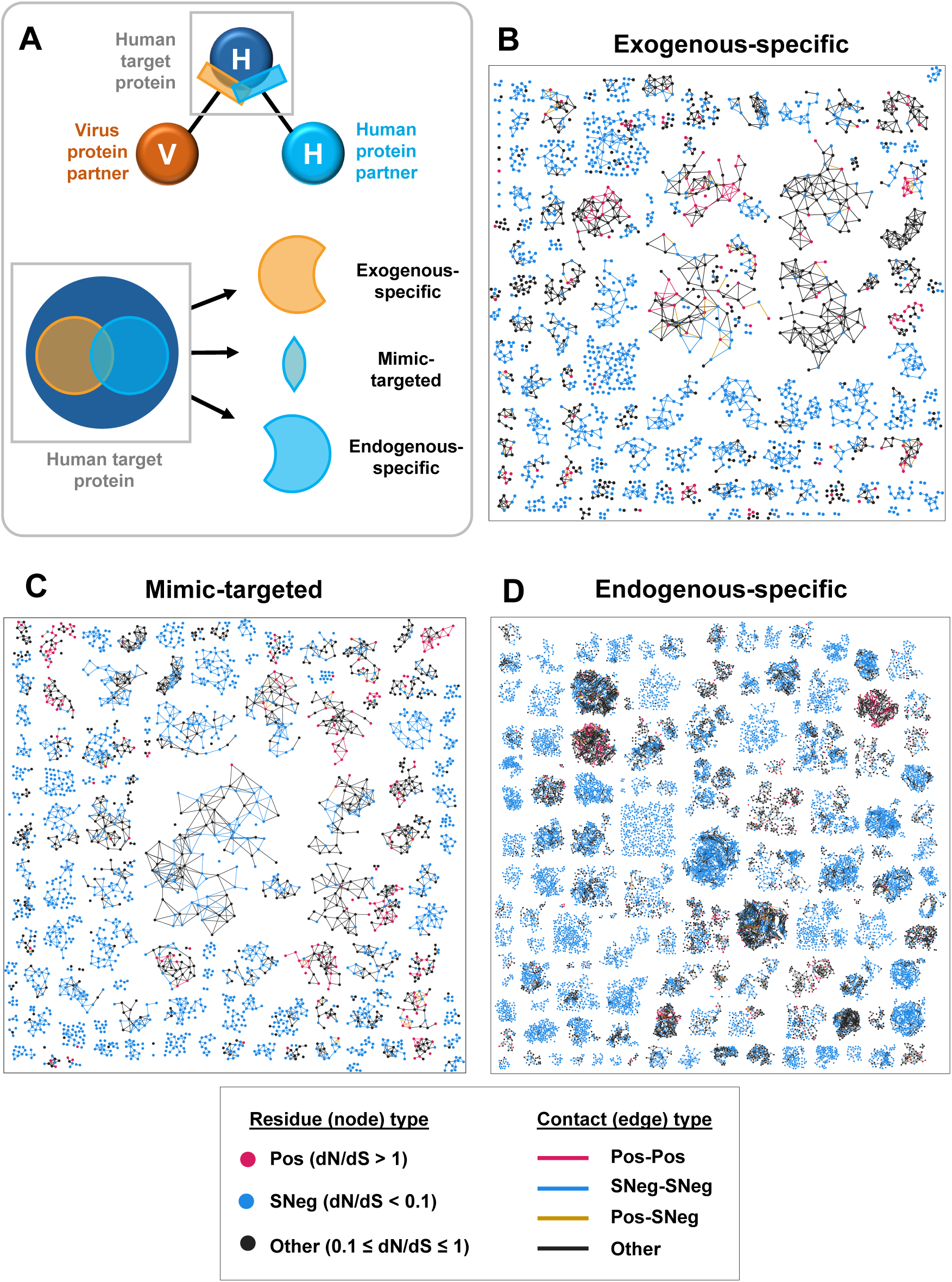
Sub-interface categorization and sub-interface-specific contact graphs. **(A)** Top: Diagram depicting a human target protein being targeted by both a viral protein partner and a human protein partner at overlapping PPI interfaces. Bottom: Categorizing virus-targeted (all exogenous, orange circle) and human-targeted (all endogenous, light blue circle) PPI interfaces on a human target protein (dark blue circle) into three sub-interfaces: (i) exogenous-specific (virus-targeted only), (ii) mimic-targeted (both virus-targeted and human-targeted), and (iii) endogenous-specific (human-targeted only). **(B)-(D)** Residue-residue contact graphs for sub-interfaces (exogenous-specific, mimic-targeted, and endogenous-specific) on human target proteins (see **Materials and Methods** for details on constructing sub-interface contact graphs). Each cluster in the contact graph represents the corresponding sub-interface of an individual human target protein. Nodes represent interfacial residues within that sub-interface on human target proteins and edges represent a pair of interfacial residues in contact. Nodes and edges follow the same labeling scheme as in the original contact graph (Figure 1A). Contact graphs were visualized using Graphviz [63].

The contact enrichment and depletion trends for the entire exogenous interface (all exogenous) and the entire endogenous interface (all endogenous) contact graphs (Figures 4A and **4B**) are qualitatively consistent with the original contact graph involving all interfacial residues ( Figure 1B). Namely, both all exogenous and all endogenous interface contact graphs exhibit high enrichment of Pos-Pos contacts, very little to no enrichment of SNeg-SNeg contacts, strong depletion of Pos-SNeg contacts, and little to no enrichment or depletion of all other contacts.

**Figure 4.**
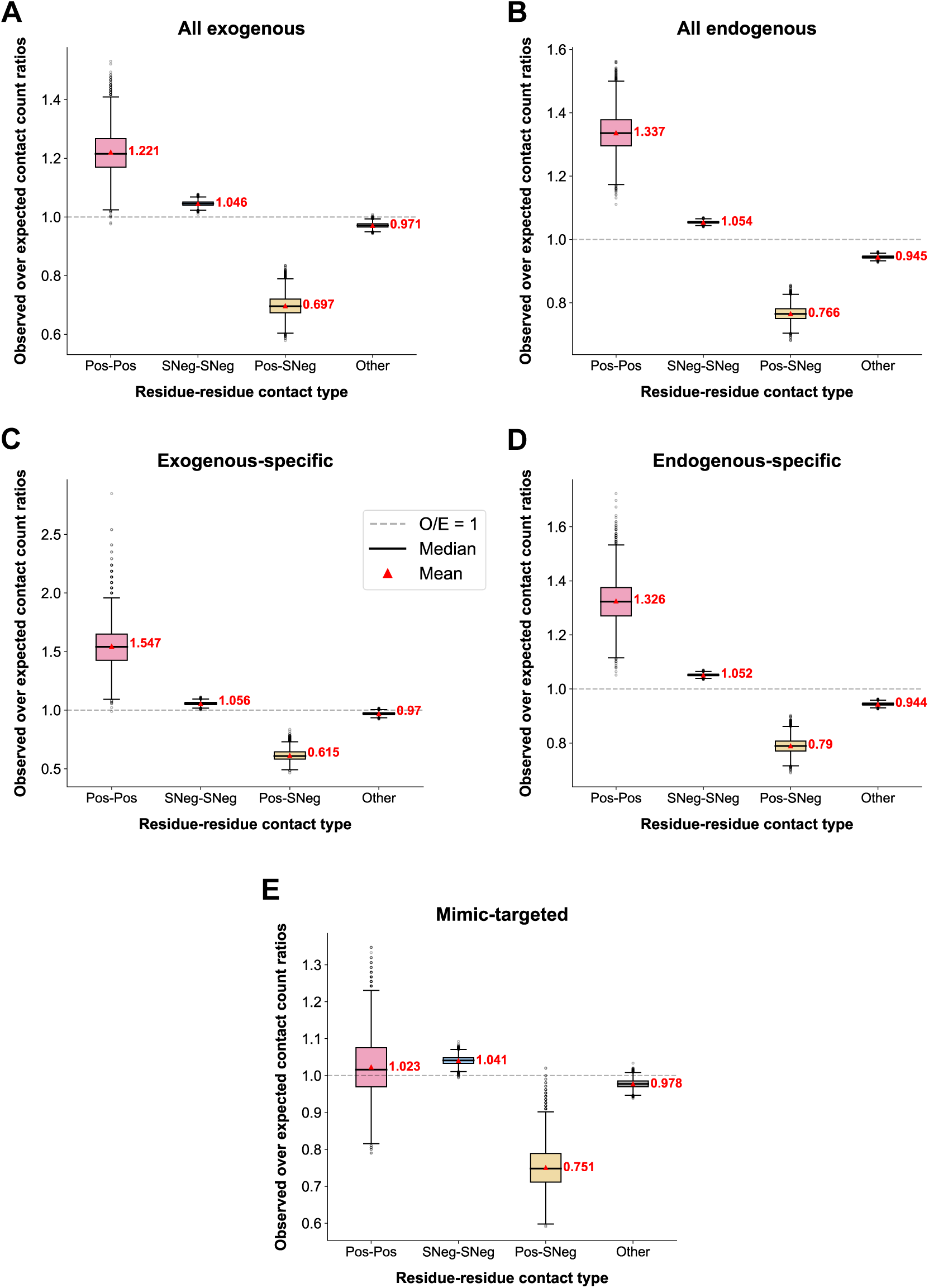
O/E contact count ratios for interface and sub-interface contact graphs. **(A)-(E)** Boxplots illustrating the distribution of observed-over-expected (O/E) contact counts for different residue-residue contact types, based on 10,000 intra-protein node-label randomization trials, for interface and sub-interface contact graphs: **(A)** all exogenous interface (exogenous-specific and mimic-targeted combined), **(B)** all endogenous interface (endogenous-specific and mimic-targeted combined), **(C)** exogenous-specific sub-interface, **(D)** endogenous-specific sub-interface, and **(E)** mimic-targeted sub-interface. The gray dashed line represents O/E = 1. Overall (mean) O/E ratios (across the randomization trials) are represented by red triangles and labeled in red, while median O/E ratios (across the randomization trials) are represented by black horizontal lines. Since all overall O/E ratios are significantly different from 1 (*p* < 0.001, two-tailed one-sample t-test against 1), we focus our analysis on effect size rather than statistical significance.

Interestingly, when we split the entire exogenous and entire endogenous interfaces into their separate sub-interfaces (exogenous-specific, endogenous-specific, and mimic-targeted; Figure 3), we observe that while the contact enrichment and depletion trends for exogenous-specific (Figure 4C) and endogenous-specific (Figure 4D) contact graphs are also qualitatively consistent with the original all interfacial residues contact graph (Figure 1B), the mimic-targeted contact graph (Figure 4E) displays trends that are distinct from all the other interfaces. Specifically, although we observe high enrichment of Pos-Pos contacts in exogenous-specific and endogenous-specific sub-interfaces, we observe very little to no enrichment of Pos-Pos contacts within the mimic-targeted sub-interface. Such findings suggest that while positively selected residues are spatially clustered on exogenous-specific and endogenous-specific sub-interfaces, they are more uniformly distributed across mimic-targeted sub-interfaces. Moreover, the fact that Pos-SNeg contacts remain depleted in mimic-targeted sub-interfaces (Figure 4E), where neither Pos nor SNeg residues are clustered, further enforces the notion that the spatial segregation between positive and strong negative selective pressures cannot be explained by clustering of positive selection signatures.

The observed difference in Pos-Pos contact enrichment between exogenous-specific (Figure 4C) and mimic-targeted sub-interfaces (Figure 4E) is particularly striking because in our previous work [24], we saw that the two sub-interfaces evolve at comparable rates and contain almost equal proportions of positively selected residues (0.078 for exogenous-specific and 0.081 for mimic-targeted) (Figure 5). Hence, although exogenous-specific and mimic-targeted sub-interfaces exhibit similar overall evolutionary rates and contain comparable proportions of positively selected residues, positive selection acts on only localized subsets of the exogenous-specific sub-interface, whereas it acts broadly across the entire mimic-targeted sub-interface. Such findings suggest that the host-virus arms race is centered at the mimic-targeted sub-interface. Moreover, the three sub-interface types (exogenous-specific, mimic-targeted, and endogenous-specific) can be distinguished only when both overall evolutionary rates and the spatial organization of selective pressures are considered, highlighting the importance of using both features for effective characterization of host-virus PPI interfaces.

**Figure 5.**
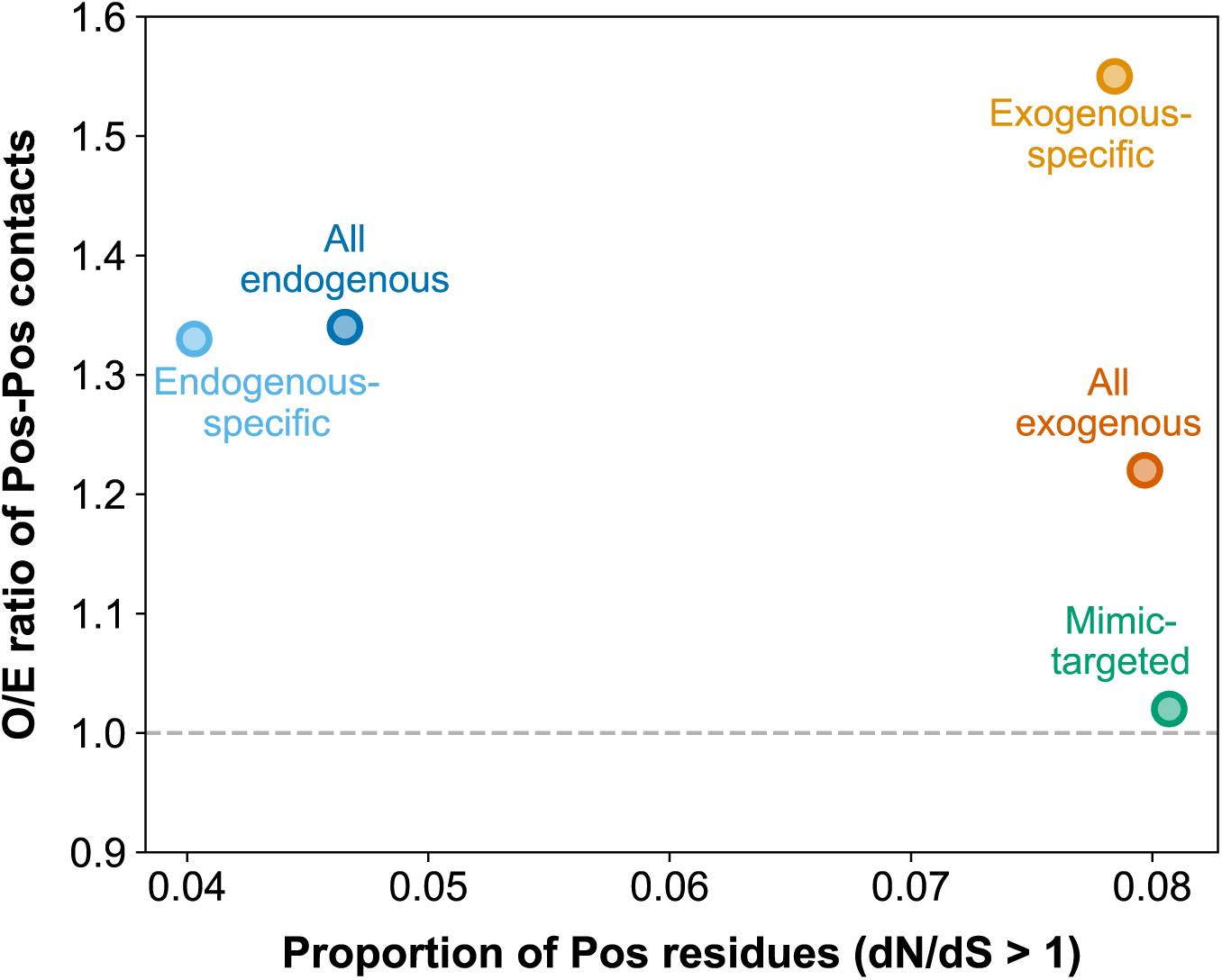
Overall O/E ratios for Pos-Pos contacts plotted against the proportion of Pos residues in each interface and sub-interface. Scatterplot illustrating the overall O/E ratio for Pos-Pos contacts (enrichment/depletion of Pos-Pos contacts) in an interface/sub-interface against the proportion of Pos residues in that interface/sub-interface. O/E = 1 is represented by the gray horizontal dashed line. Exogenous-specific and mimic-targeted sub-interfaces share similar proportions of Pos residues, but their Pos-Pos contact enrichments (and hence their spatial distribution of Pos residues) differ greatly, demonstrating that both factors are needed for effectively distinguishing distinct sub-interfaces on virus-targeted host proteins.

### Cross-interface clustering of positively selected residues is greater than within-interface clustering

To further investigate the spatial organization of positive selection signatures, we asked whether positively selected residues in the mimic-targeted interface influence clustering in exogenous-and endogenous-specific interfaces (i.e., whether Pos-Pos contacts are enriched between mimic-targeted and exogenous-specific and between mimic-targeted and endogenous-specific interfaces). We first categorized each Pos-Pos contact in the entire exogenous interface into three location categories: (i) “within exogenous-specific”, (ii) “between exogenous-specific and mimic-targeted”, and (iii) “within mimic-targeted”. Similarly, we categorized each Pos-Pos contact in the entire endogenous interface into three location categories: (i) “within endogenous-specific”, (ii) “between endogenous-specific and mimic-targeted”, and (iii) “within mimic-targeted”. We then calculated O/E ratios for each location category by comparing observed Pos-Pos contacts to expectations derived from intra-protein randomization of residue labels within each interface separately, ensuring that the interface-specific proportions of positively selected residues were preserved.

Cross-interface O/E values reveal strong clustering of positive selection between the mimic-targeted and exogenous-specific interfaces (Figure 6A, O/E = 1.752) and between the mimic-targeted and endogenous-specific interfaces (Figure 6B, O/E = 1.544), whereas no Pos-Pos contacts were observed between the exogenous-and endogenous-specific interfaces. Within-interface O/E ratios remain broadly consistent with our previous analysis (Figure 4), showing clustering of positively selected residues within the exogenous-and endogenous-specific interfaces and a more uniform distribution of positively selected residues within the mimic-targeted interface. Notably, clustering of positive selection between mimic-targeted and exogenous-specific interfaces is significantly greater than that observed within the exogenous-specific interface itself (Figure 6A), and clustering of positive selection between mimic-targeted and endogenous-specific interfaces is significantly greater than that observed within the endogenous-specific interface itself (Figure 6B) (two-tailed Mann-Whitney U test, *p* < 0.001 for both comparisons). The broad distribution of positively selected residues across the mimic-targeted interface, together with significantly greater enrichment of Pos-Pos contacts spanning into both exogenous-and endogenous-specific interfaces compared with within-interface clustering, suggests that the mimic-targeted interface is a focal point of virus-driven adaptation.

**Figure 6.**
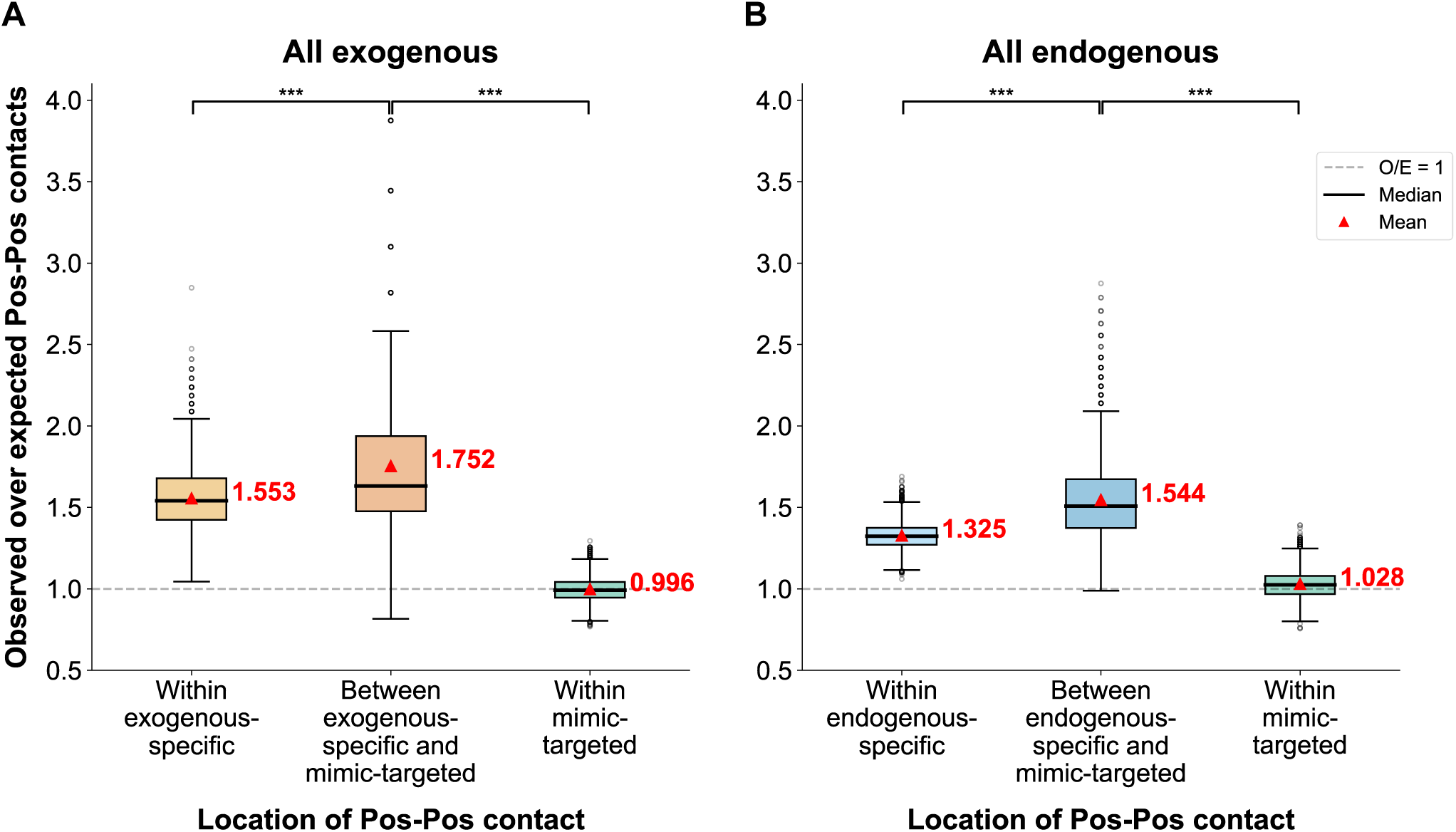
Pos-Pos contact enrichments at different locations across the entire virus-targeted (all exogenous) and entire endogenous-targeted (all endogenous) interfaces **(A)** Boxplots illustrating the distribution of O/E contact counts for Pos-Pos contacts at different locations in the entire virus-targeted (all exogenous) interface: (i) within the exogenous-specific sub-interface, (ii) between exogenous-specific and mimic-targeted sub-interfaces, and (iii) within the mimic-targeted sub-interface. **(B)** Boxplots illustrating the distribution of O/E contact counts for Pos-Pos contacts at different locations in the entire endogenous-targeted (all endogenous) interface: (i) within the endogenous-specific sub-interface, (ii) between endogenous-specific and mimic-targeted sub-interfaces, and (iii) within the mimic-targeted sub-interface. **(A)-(B)** Distributions of O/E ratios are based on 10,000 within-protein node-label randomization trials. The gray dashed line represents O/E = 1. Overall (mean) O/E ratios (across the randomization trials) are represented by red triangles and labeled in red, while median O/E ratios (across the randomization trials) are represented by black horizontal lines. A two-tailed Mann-Whitney U test was used to assess whether the distribution of O/E values in one location differed significantly from that in another location. Statistical significance is denoted as follows: \*\*\**p* < 0.001.

## Discussion

By overlaying spatial selection patterns onto an evolutionarily annotated high-resolution map of human-virus protein-protein interaction interfaces, we demonstrated that selective pressures are heterogeneously distributed across the interfaces of virus-targeted host proteins, with this heterogeneity observed both between selection types and across different interface contexts.

At the selection level, positive and negative selection exhibit distinct spatial organizations across host PPI interfaces collectively (Figure 1B). The clustering of positively selected residues observed globally across PPI interfaces is broadly consistent with previous studies showing that spatial clustering of substitutions in protein structures is associated with functionally adaptive changes [47], and that positively selected sites form clusters and are situated in regions of functional importance [25]. While those analyses were conducted at the scale of entire protein structures, our interface-focused perspective demonstrates that similar spatial organization is evident broadly across host PPI interfaces, reinforcing the idea that adaptive evolution is structurally localized rather than randomly distributed. However, by further partitioning host PPI interfaces based on interaction context and examining spatial selection patterns at this finer scale (Figures 3 **and 4**), we find that positive selection clustering is not universally observed across all interface types, indicating that the arrangement of adaptive residues depends on interaction context. These findings suggest that individual interface-specific patterns of adaptive change are more heterogeneous than can be observed through protein-wide or global interface-wide analyses.

Additionally, we find that positively and strongly negatively selected residues are spatially segregated across host-virus PPI interfaces (Figure 1B), reflecting a functional partitioning of adaptive and strongly constrained regions. We note that weaker negatively selected residues do not show the same segregation, suggesting that the observed separation primarily involves the most strongly constrained sites. This pattern may indicate that adaptation and strong constraint act largely independently, and potentially antagonistically, with adaptive residues concentrated in regions less constrained by essential structural or functional roles. Such an arrangement could allow host proteins to accommodate adaptive changes in response to viral pressures while maintaining critical interaction sites, hinting at a modular organization in which adaptive and constrained regions occupy complementary structural niches.

At the interface level, virus-targeted interfaces themselves can be characterized by the patterns of positive selection they exhibit (Figures 4C **and 4E**). Despite sharing similar proportions of positively selected residues, exogenous-specific interfaces were found to display clusters of positive selection, whereas mimic-targeted interfaces showed a more uniform distribution of positive selection (Figure 5). This suggests that the spatial distribution of positively selected residues varies independently from the overall proportion of positively selected residues in an interface. Moreover, the difference in spatial organization of positive selection across exogenous-specific and mimic-targeted interfaces reflects distinct evolutionary strategies. Clustered positive selection in exogenous-specific interfaces suggests focused adaptation to specific viral contacts, whereas dispersed positive selection in mimic-targeted interfaces suggests a strategy of distributed adaptation, whereby changes are spread across the interface to support flexible responses to viral targeting while preserving functional integrity and minimizing vulnerability at any single site. Similar patterns have been suggested for protein kinase R (PKR), a host protein in which viral mimics such as vaccinia virus K3L compete with its substrate eIF2α for binding at the kinase domain [17,19,48]. Evolutionary analyses of PKR’s kinase domain indicate that positively selected residues occur across several regions (including sites mediating eIF2α binding) rather than being concentrated at a single site, consistent with the idea that distributed adaptation could help PKR escape K3L exploitation while preserving eIF2α binding [17,19]. Our observation of broadly dispersed positive selection signatures across mimic-targeted interfaces, based on the full residue-residue contact graph spanning multiple human target proteins, suggests that distributed adaptation at mimic-targeted interfaces may represent a general host strategy to counteract viral mimicry, rather than a feature unique to a single protein.

Our observation that positively selected residues cluster between distinct PPI interfaces further highlights mimic-targeted interfaces as focal points of adaptive pressure. In mimic-targeted interfaces, positively selected residues are broadly distributed across the entire interface but appear to influence clustering across its interface boundaries with both exogenous-specific and endogenous-specific interfaces (Figure 6). Specifically, cross-interface positive selection clustering between mimic-targeted and exogenous-specific residues, as well as between mimic-targeted and endogenous-specific residues, exceed the clustering observed within exogenous-specific or endogenous-specific interfaces alone. This suggests that the broadly distributed positive selection signatures across mimic-targeted interfaces may serve as an anchor for coordinating adaptive changes across neighboring interfaces. Such observations also point toward a potential gradient effect, in which adaptive changes at mimic-targeted interfaces could propagate to more distal interface regions, with their influence diminishing with distance from the mimic-targeted interface. While long-range propagation has been documented for residues under negative selection in the context of catalytic residues [32], our findings raise the possibility that a similar principle may apply for positive selection in the context of host-virus PPI interfaces. Formal testing of this gradient hypothesis will require larger datasets or proteins with more extensive interaction interfaces, as the inherently small sizes of virus-host PPI interfaces [49], their further subdivision into distinct sub-interfaces, and the sparsity of positively selected residues [50,51] constrain the number of sites available for robust analysis. Nonetheless, our findings provide a conceptual foundation for understanding how adaptive pressure is organized across different host-virus PPI interfaces and position mimic-targeted interfaces as central regions for future studies of host-virus evolutionary dynamics.

Together, these insights elucidate how host proteins reconcile the preservation of endogenous functions with adaptation to viral binding. By examining spatial patterns both across different types of selective pressures and across distinct interfaces on virus-targeted host proteins, our study reveals a context-dependent and coordinated organization of selective pressures, highlighting the precise strategies hosts employ to navigate their evolutionary landscape in the face of viral antagonists.

## Materials and Methods

### Identification of interfacial residues and estimation of residue-level evolutionary rates

Interfacial residues and residue-level evolutionary rates were defined following previously described procedures [24], specified as follows. Exogenous (virus-human) and endogenous (within-human) protein-protein interactions (PPIs) from the IntAct [52] database were mapped onto three-dimensional structures using either experimentally determined complexes from the Protein Data Bank (PDB) [33] or homology-based structural templates (constructed via sequence alignment of interacting proteins to subunits in PDB protein complexes). Interfacial residues (i.e., residues that are in physical contact with an interaction partner) on virus-targeted human proteins (referred to as human target proteins) were defined as residues whose relative solvent accessibility decreased upon binding to an interaction partner.

Residue-level evolutionary rates (site-specific dN/dS values) for interfacial residues were estimated using HyPhy’s one-rate Fixed Effects Likelihood (FEL) method [53,54], with human target proteins as the reference and orthologs from 19 other mammals used for comparative analyses. These orthologs were identified via reciprocal best-hit amino acid searches conducted between the human proteome and each of the 19 other mammalian proteomes. Coding sequences for both human target proteins and their orthologs were retrieved from RefSeq [55] and Ensembl [56], with tBLASTn used to select the best-matching transcripts. Amino acid sequences translated from these transcripts were aligned using MAFFT [57], and the resulting amino acid alignments were used to guide codon alignments with PAL2NAL [58]. A phylogenetic species tree of the 20 species, obtained from the TimeTree database [59], was used to guide rate estimation. Site-specific dN/dS values were then calculated for each codon position (following the protocol described in Sydykova et al. [60]) and mapped back to the human protein sequence.

In contrast to prior work, which summarized evolutionary rates at the interface level (e.g., interface averages and threshold-based fractions), the present study provides the first residue-level analysis, retaining individual site-specific dN/dS values for each interfacial residue and integrating them into residue-residue contact graphs to examine the spatial organization of selective pressures across host PPI interfaces.

All subsequent analyses in this study, such as constructing residue-residue contact graphs and assessing the spatial distribution of selective pressures, use these interfacial residues and site-specific dN/dS values as input and represent new methodological components introduced here.

### Residue-residue distance calculations and contact definition

For each human target protein, we calculated the distance between every pair of interfacial residues within that protein. The distance between a pair of residues (i.e., residue-residue distance) was defined as the minimum Euclidean distance between any pair of atoms between the two residues, based on the highest-resolution structural template (obtained from the above structural characterization procedure) that contains both residues. Two residues were considered to be in contact if their residue-residue distance was less than 5 Å.

### Constructing a residue-residue contact graph of interfacial residues within the structural human-virus protein-protein interaction network

We constructed a residue-residue contact graph by treating each interfacial residue on human target proteins as a node and adding an edge between every pair of residues in contact (as defined above). Within the contact graph, nodes (residues) were colored based on their corresponding site-specific dN/dS values: (i) red for dN/dS > 1 (positive selection, “Pos”), (ii) blue for dN/dS < 0.1 (strong negative selection, “SNeg”), and (iii) black for 0.1 ≤ dN/dS ≤ 1 (all remaining residues, “Other”). In this study, we used a more stringent negative selection threshold (dN/dS < 0.1), rather than the conventional dN/dS < 1 threshold [61,62], to ensure a more balanced comparison with positive selection (dN/dS > 1). Edges (residue-residue contacts) were colored based on the contact type: (i) red: two positively selected residues in contact (denoted “Pos-Pos”), (ii) blue: two strongly negatively selected residues in contact (denoted “SNeg-SNeg”), (iii) amber: a positively selected residue and a strongly negatively selected residue in contact (denoted “Pos-SNeg”), and (iv) black: all other pairs of residues in contact (denoted “Other”), which contains three subcategories (“Pos-Other”, “SNeg-Other”, and “Other-Other”). Each cluster (subgraph) within the contact graph represents an individual human target protein.

### Whole contact graph residue-residue contact count enrichment analysis

For each residue-residue contact type (as defined in the above section), we computed the observed-over-expected (O/E) occurrence of that contact type. The observed occurrence of a contact type (O) corresponds to the number of edges corresponding to that contact type in the entire contact graph (constructed above). To obtain the expected occurrence (E) of a given contact type across the entire contact graph, we performed intra-protein node-label (“Pos”, “SNeg”, and “Other”) permutations to create a randomized contact graph (i.e., node labels within the original contact graph were shuffled within each protein separately while edges remained intact). We conducted 10,000 iterations of this randomization approach. To derive an overall representative measure of enrichment (or depletion) of a contact type, we plotted the distribution of O/E ratios across the 10,000 permutation trials (where O remains fixed in each trial), and reported the overall (mean) O/E ratio based on these trials. The overall O/E ratio was assessed as follows: (i) overall O/E >> 1: enrichment of a contact type, (ii) overall O/E ≈ 1: little to no enrichment or depletion of a contact type, (iii) overall O/E << 1: depletion of a contact type. In this work, effect sizes were used as the primary measure of enrichment rather than statistical significance, as the majority of overall O/E ratios are significantly different from 1 (*p* < 0.001, two-tailed one-sample t-test against 1).

### Per-protein residue-residue contact count enrichment analysis

To derive contact count enrichments on a per-protein basis, we split a contact graph into its individual protein subgraphs and performed the same procedure as above for one protein subgraph at a time. This resulted in an individual overall (mean) O/E ratio (calculated across 10,000 node-label permutation trials) for each protein. To obtain a whole-graph measure of contact count enrichment, we plotted the distribution of the per-protein overall O/E ratios, and reported the mean and median of these per-protein overall O/E ratios. To ensure that this per-protein analysis was conducted on sufficiently large and reliable data, we excluded protein subgraphs with limited Pos or SNeg residues (i.e., less than 5 Pos or less than 5 SNeg residues).

### Constructing contact graphs for specific interfaces and sub-interfaces

Interfacial residues on human target proteins were categorized into two interfaces: (i) all exogenous (targeted by viral proteins), and (ii) all endogenous (targeted by human proteins). These two interfaces were further categorized into three sub-interfaces (Figure 3A): (i) exogenous-specific (targeted only by viral proteins), (ii) mimic-targeted (targeted by both viral and human proteins), and (iii) endogenous-specific (targeted only by human proteins). To obtain contact graphs for specific interfaces (all exogenous and all endogenous) or sub-interfaces (exogenous-specific, mimic-targeted, and endogenous-specific), we constructed subgraphs from residues corresponding to the interface or sub-interface of interest, retaining only residue-residue contacts between them that existed in the original (all interfacial residues) contact graph. We used the same procedure as for the original contact graph to assess enrichment or depletion of a contact type.

### Assessing and comparing Pos-Pos contact enrichment within and between sub-interfaces

We split Pos-Pos contacts in the combined exogenous (all exogenous) interface and in the combined endogenous (all endogenous) interface into three location categories: (i) those that occur within exogenous-specific (for all exogenous) or endogenous-specific (for all endogenous) sub-interfaces, (ii) those that occur across sub-interface boundaries (i.e., between exogenous-specific and mimic-targeted interfaces for all exogenous, and between endogenous-specific and mimic-targeted interfaces for all endogenous). (iii) those that occur within the mimic-targeted sub-interface (for both all exogenous and all endogenous)

To derive the expected number of Pos-Pos contacts within or across sub-interface boundaries, we randomized node-labels by performing intra-protein node-label permutations within exogenous-specific, endogenous-specific, and mimic-targeted sub-interfaces separately to preserve original proportions of distinct selective pressures within the individual sub-interfaces. We performed 10,000 iterations of this randomization approach and plotted boxplots to illustrate the distribution of observed over expected (O/E) contact count ratios across the 10,000 randomization trials and reported the overall (mean) O/E ratio. To compare overall O/E ratios in different locations (within and across sub-interface boundaries), we used a two-tailed Mann-Whitney U test to assess whether the distribution of O/E values across the randomization trials in one location differed significantly from that in another location.

## Data and code availability

All data and code used to conduct this study is available in the following GitHub repository: https://github.com/wanchunsu/SpatiOEvo.

## Acknowledgements

This work was supported by Natural Sciences and Engineering Research Council of Canada grants RGPIN-2019-05952 and RGPAS-2019-00012, Canada Foundation for Innovation grants JELF-33732 and IF-33122, and Canada Research Chairs program (CRC-2022-00424). W.-C.S. is supported by a doctoral fellowship from Fonds de recherche du Québec – Nature et technologies.

