## Supplementary Figures S1-S7, Table S1 for "Divergent landscapes of positive and negative selection signatures across residue-resolved human-virus protein-protein interaction interfaces"

### Supplementary Materials

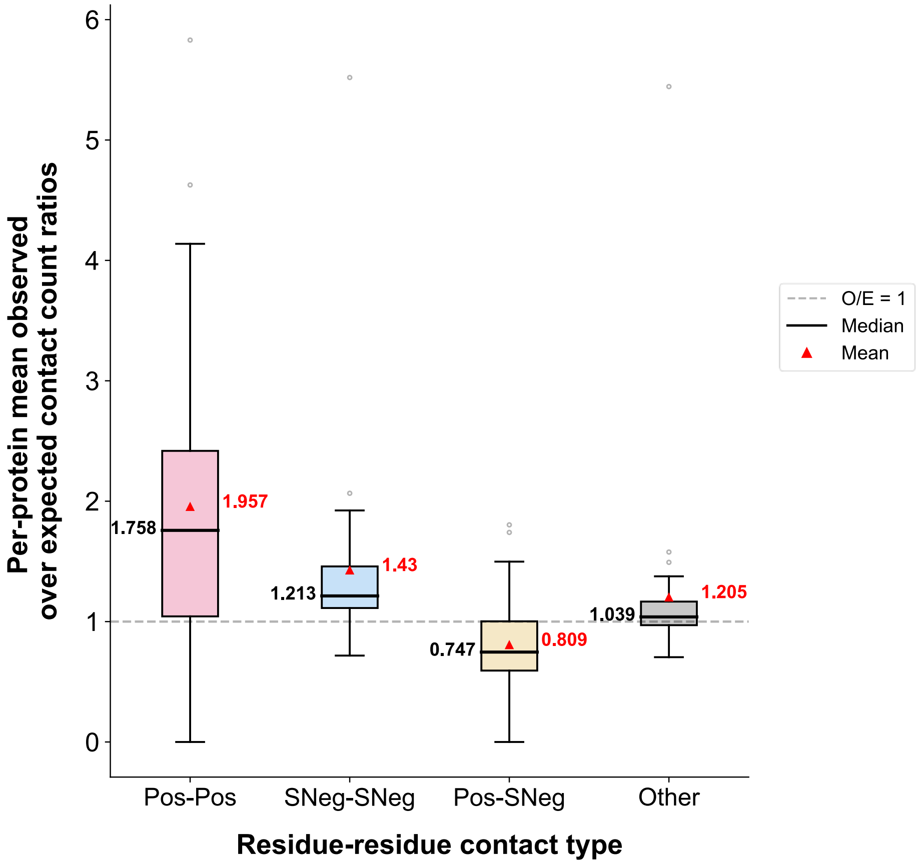


**Figure S1. Per-protein O/E ratios.** Each box depicts the distribution of per-protein mean (overall) O/E ratios (based on 10,000 permutation trials), with one datapoint per protein, for a specific residue-residue contact type. Proteins with limited Pos or SNeg residues (i.e., less than 5 Pos residues or less than 5 SNeg residues) are not included in this plot. Mean and median values across the distribution of per-protein overall O/E ratios are depicted in red (triangles and labels) and black (horizontal lines and labels), respectively.

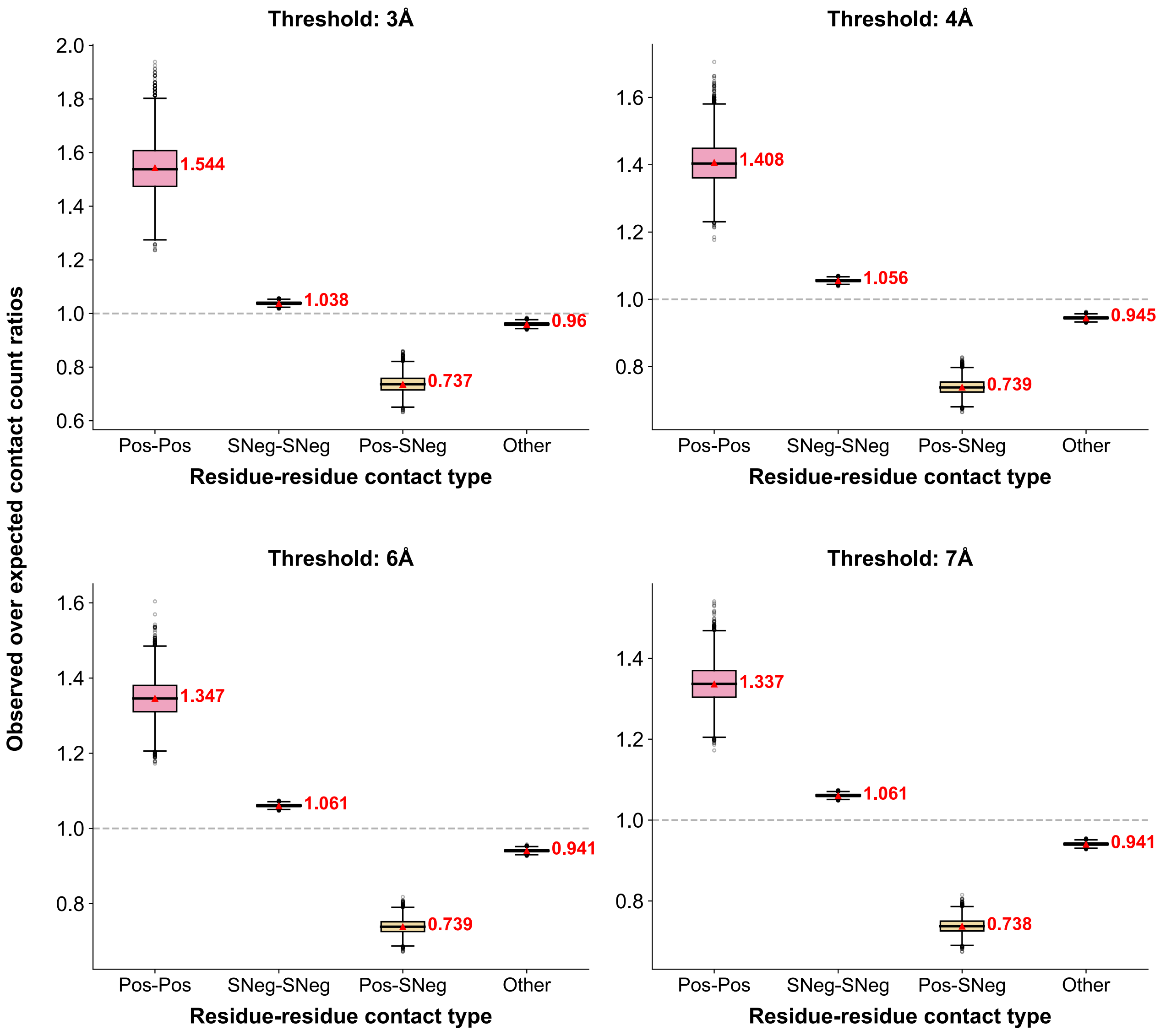


**Figure S2. Altering the residue-residue contact threshold and resulting O/E contact count ratios.** Enrichment/depletion trends are qualitatively consistent with the original analysis (threshold: 5Å).

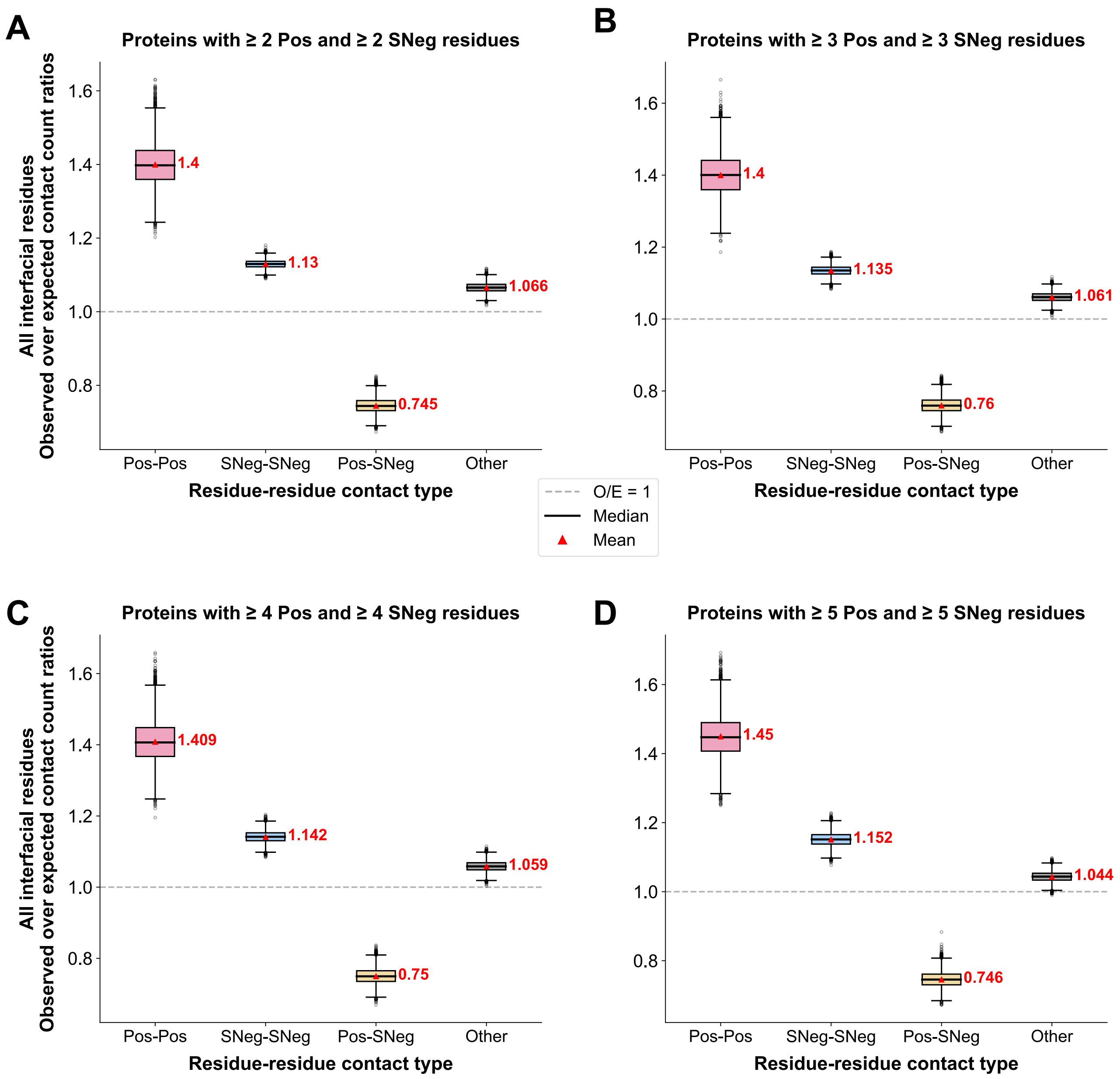


**Figure S3. Observed-over-expected contact count ratios for different subgraphs comprising proteins with sufficient numbers of Pos and SNeg residues.**

**(A)-(D)** Boxplots illustrating the distribution of observed-over-expected (O/E) contact counts for different residue-residue contact types, based on 10,000 intra-protein node-label randomization trials. Each plot corresponds to a contact graph that includes only human target proteins with at least n Pos and n SNeg interfacial residues, where n = 2, 3, 4, and 5 for subfigures A to D, respectively. The gray dashed line represents O/E = 1. Overall (mean) O/E ratios (across the randomization trials) are represented by red triangles and labeled in red, while median O/E ratios (across the randomization trials) are represented by black horizontal lines. Since all overall O/E ratios are significantly different from 1 ($p < 0.001$, two-tailed one-sample t-test against 1), we focus our analysis on effect size rather than statistical significance. Together, these plots illustrate the robust segregation between positive selection and strong negative selection (Pos-SNeg contacts remain depleted to a similar extent as in **Figure 1B**), as well as consistent enrichment/depletion trends for all other contact types.

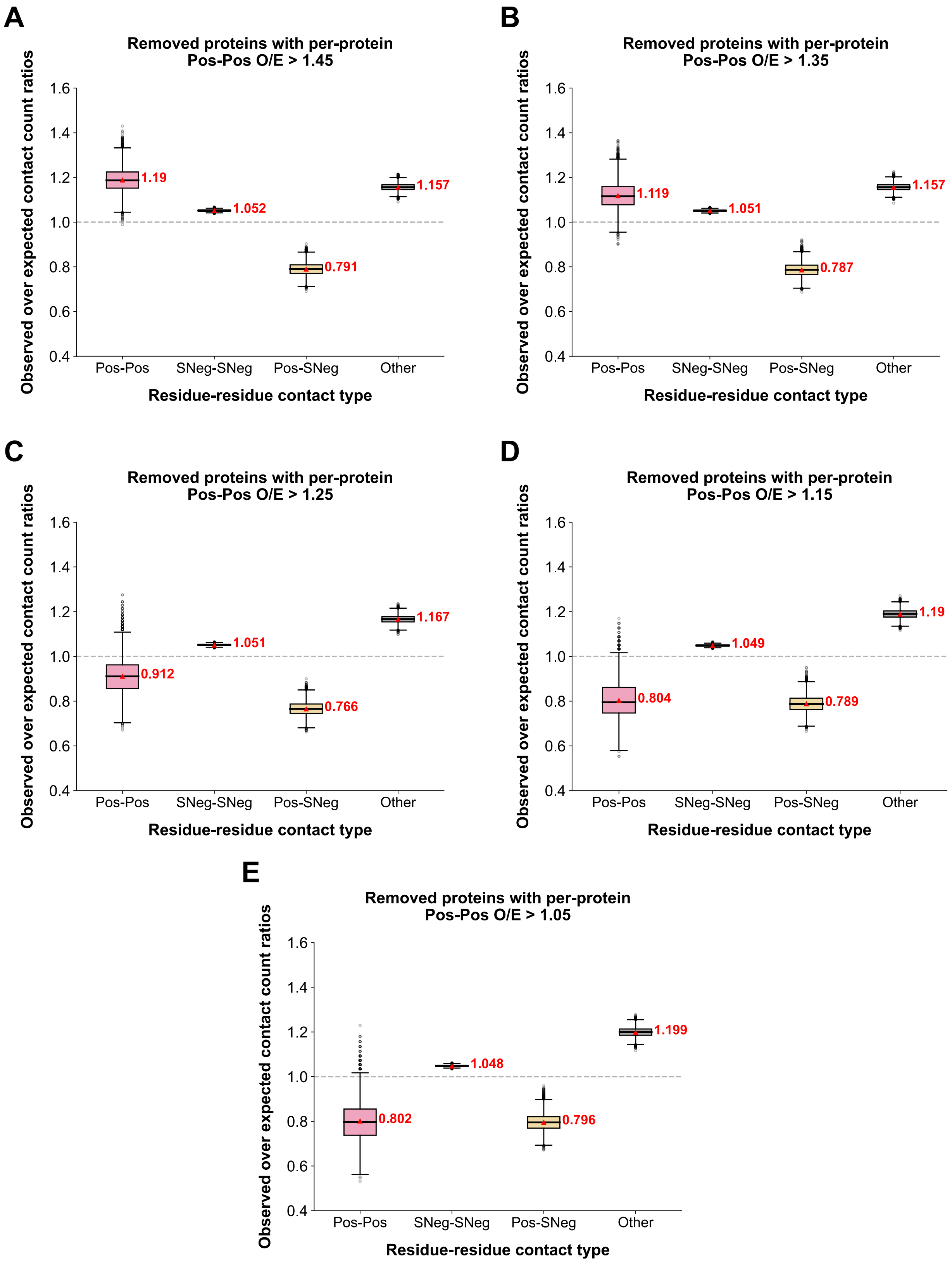


**Figure S4. Removing proteins with varying degrees of Pos clustering.** Boxplots depicting the O/E contact count ratios across the entire contact graph after removal of proteins with per-protein Pos-Pos O/E ratios greater than a given threshold: **(A)** O/E > 1.45, **(B)** O/E > 1.35, **(C)** O/E > 1.25, **(D)** O/E > 1.15, and **(E)** O/E > 1.05.

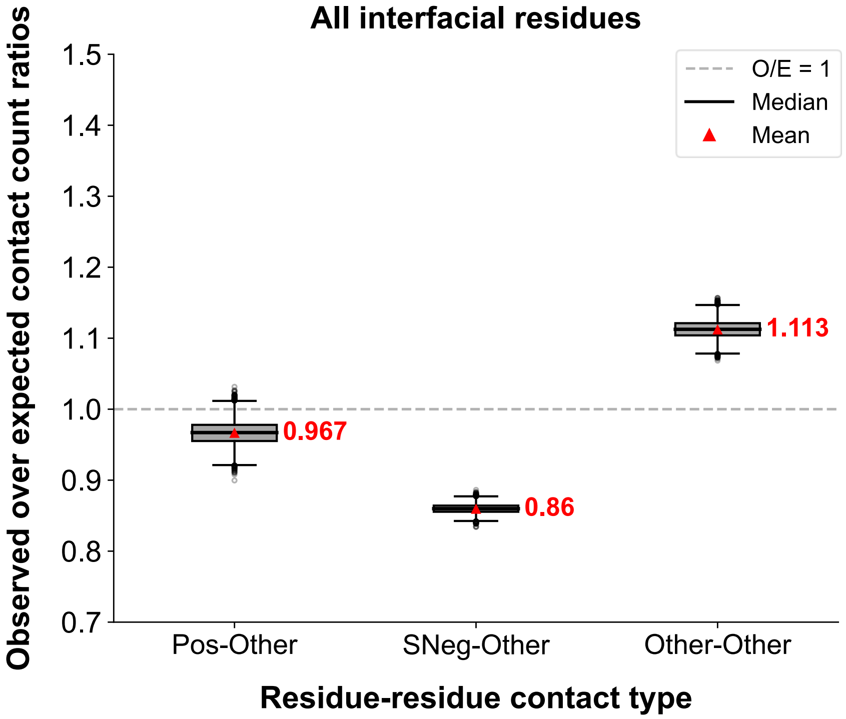


**Figure S5. O/E contact count ratios for contacts involving "Other" (0.1 ≤ dN/dS ≤ 1) residues.** Compared to Pos-Pos and Pos-SNeg (**Figure 1B**), the three contact types here exhibit very small effect sizes (O/E ≈ 1).

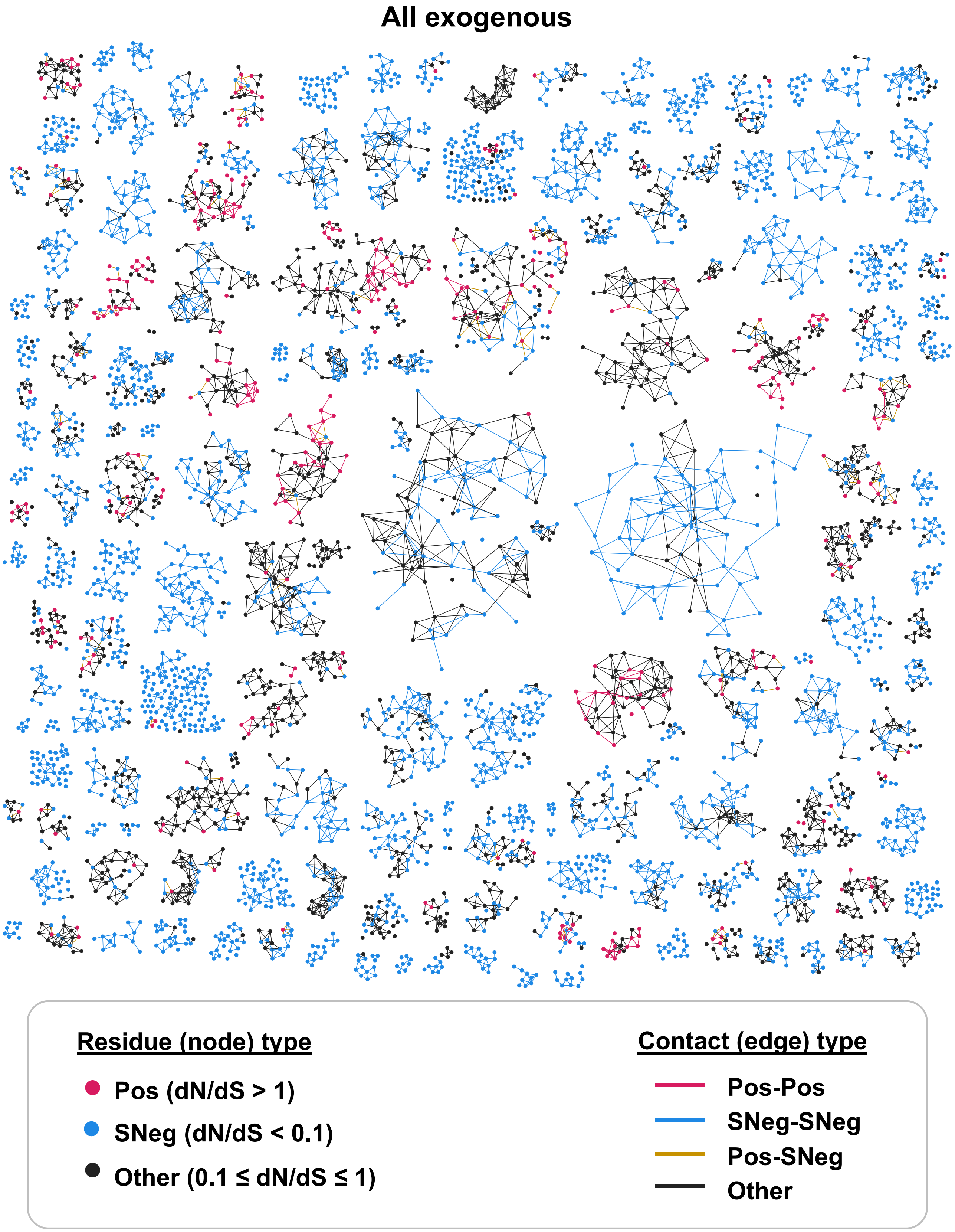


**Figure S6. Residue-residue contact graph of all virus-targeted interfaces (all exogenous) on human target proteins.**

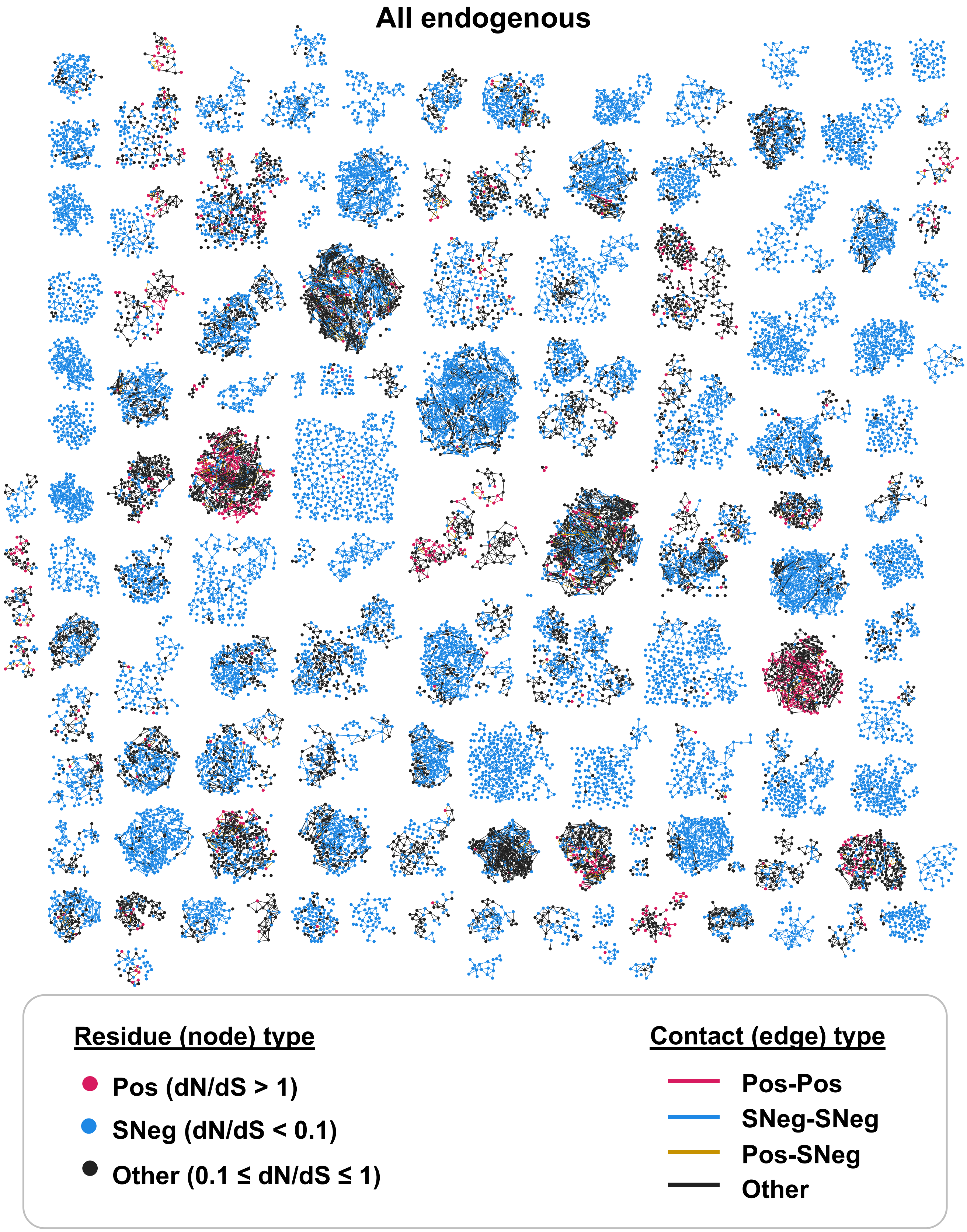


**Figure S7. Residue-residue contact graph of all within-host interfaces (all endogenous) on human target proteins.**

**Table S1. Number of human target proteins and number of edges (for each contact type) in the whole interface and interface-specific contact graphs.**

| | Human target proteins | Pos-Pos contacts | SNeg-SNeg contacts | Pos-SNeg contacts | Other contacts |
|---|---|---|---|---|---|
| **Whole interface** | 159 | 702 | 21,435 | 898 | 17,518 |
| **All exogenous** | 157 | 248 | 3,622 | 206 | 3,778 |
| **All endogenous** | 132 | 561 | 19,455 | 778 | 15,421 |
| **Exogenous-specific** | 115 | 94 | 1,320 | 92 | 1,446 |
| **Endogenous-specific** | 129 | 348 | 15,755 | 591 | 11,937 |
| **Mimic-targeted** | 118 | 128 | 1,952 | 101 | 1,963 |
